# Mitochondrial Morphology Regulates Organellar Ca^2+^ Uptake and Changes Cellular Ca^2+^ Homeostasis

**DOI:** 10.1101/624981

**Authors:** Alicia J. Kowaltowski, Sergio L. Menezes‐Filho, Essam Assali, Isabela G. Gonçalves, Phablo Abreu, Nathanael Miller, Patricia Nolasco, Francisco R. M. Laurindo, Alexandre Bruni‐Cardoso, Orian Shirihai

## Abstract

Changes in mitochondrial size and shape have been implicated in several physiological processes, but their role in mitochondrial Ca^2+^ uptake regulation and overall cellular Ca^2+^ homeostasis is largely unknown. Here we show that modulating mitochondrial dynamics towards increased fusion through expression of a dominant negative form of the fission protein DRP1 (DRP1‐DN) markedly increased both mitochondrial Ca^2+^ retention capacity and Ca^2+^ uptake rates in permeabilized C2C12 cells. Similar results were seen using the pharmacological fusion‐promoting M1 molecule. Conversely, promoting a fission phenotype through the knockdown of the fusion protein mitofusin 2 (MFN2) strongly reduced mitochondrial Ca^2+^ uptake speed and capacity in these cells. These changes were not dependent on modifications in inner membrane potentials or the mitochondrial permeability transition. Implications of mitochondrial morphology modulation on cellular calcium homeostasis were measured in intact cells; mitochondrial fission promoted lower basal cellular calcium levels and lower endoplasmic reticulum (ER) calcium stores, as measured by depletion with thapsigargin. Indeed, mitochondrial fission was associated with ER stress. Additionally, the calcium‐replenishing process of store‐operated calcium entry (SOCE) was impaired in MFN2 knockdown cells, while DRP1‐DN‐promoted fusion resulted in faster cytosolic Ca^2+^ increase rates. Overall, our results show a novel role for mitochondrial morphology in the regulation of mitochondrial Ca^2+^ uptake, which impacts on cellular Ca^2+^ homeostasis.

## Introduction

Recent advances in imaging techniques have allowed for detailed studies of mitochondrial morphology and their dynamic changes in live cells. Mitochondria not only vary in size and shape in different cell types, but also rapidly remodel their morphology in response to environmental changes such as nutrient availability (Chan, 2006; Westerman, 2010; Liesa et al., 2009; Liesa and Shirihai, 2013; Pernas and Scorrano, 2016). Interestingly, changes in mitochondrial morphology occur not only in response to metabolic cues, but also reciprocally regulate cellular metabolic responses (Liesa and Shirihai, 2013). Dynamic changes in mitochondrial morphology additionally regulate organelle turnover, the maintenance of a healthy mitochondrial pool and the interaction between mitochondria and other organelles such as the endoplasmic reticulum (ER) and lipid droplets (Klecker et al., 2014; Stotland and Gottlieb, 2015; Pernas and Scorrano, 2016; Valm et al., 2017; Benador et al., 2019).

The core proteins responsible for modulation of mitochondrial morphology include Mitofusin 1 and Mitofusin 2 (MFN1 and MFN2), outer membrane dynamin‐related GTPase proteins that form complexes between neighboring mitochondria and mediate outer membrane fusion. This process is followed by inner membrane fusion mediated by OPA1, a GTPase also involved in cristae remodeling (Pernas and Scorrano, 2016). MFN2 also plays a key role in Ca^2+^ signaling by mediating the interaction between mitochondria and the ER (de Brito and Scorrano, 2008) and regulating mitophagy (Chen et al., 2013). Increased MFN‐mediated mitochondrial fusion is often associated with enhanced bioenergetic efficiency (Liesa and Shirihai, 2013; Forni et al., 2016). Mitochondrial fission, on the other hand, is often associated with low bioenergetic efficiency and controlled by the cytosolic dynamin‐related protein 1 (DRP1), that assembles as oligomers around the fission site. GTP hydrolysis and DRP1 superstructure constriction then promote mitochondrial fragmentation (Pernas and Scorrano, 2016). Interestingly, the location of mitochondrial outer membrane constriction is determined by ER‐mitochondrial contact sites (Friedman et al., 2011). Recent findings also suggest that inner mitochondrial membrane fission occurs at ER‐mitochondrial contact sites in response to enhanced mitochondrial Ca^2+^ uptake from ER stores, independently of DRP1 (Chakrabarti et al., 2018).

ER‐mitochondrial interactions are important not only for the regulation of mitochondrial morphology, but also for the regulation of intracellular Ca^2+^ signals (Rizzuto et al., 1998; Csordás et al., 2006; Jackson and Robinson, 2015; Wu et al., 2018). Intracellular and intramitochondrial Ca^2+^, on the other hand, are central metabolic regulators, affecting the activity of both cytosolic and mitochondrial metabolic pathways (Clapham, 2004; Gunter and Sheu, 2009; Vercesi et al., 2018). While the close interaction between the ER and mitochondria is important for adequate signaling, recent studies have shown that ER‐mitochondrial interactions increase excessively in high fat diets, resulting in mitochondrial Ca^2+^ overload and dysfunction. Disrupting these interactions can increase animal health despite the diet, demonstrating the importance of mitochondrial Ca^2+^ in metabolic control (Arruda et al., 2014; Arruda and Hotamisligil, 2015).

Ca^2+^ uptake into the mitochondrial matrix occurs through a mitochondrial Ca^2+^ uniporter (MCU) and is driven by the inner mitochondrial membrane potential (ΔΨ), which attracts positively charged species (Baughman et al., 2011). Mitochondrial Ca^2+^ uptake presents very high capacity, allowing for the accumulation of large quantities of the ion, although it is low in affinity relative to the ER. Within mitochondria, Ca^2+^ ions act as regulators of important metabolic pathways, determining ATP synthesis rates (Clapham, 2004; Gunter and Sheu, 2009; Vercesi et al., 2018). Excessive mitochondrial Ca^2+^ uptake, however, is disruptive for cell integrity under a number of pathological conditions, including stroke, ischemic heart disease and inflammatory liver conditions (Duchen, 2000; Brookes et al., 2004; Murphy and Steenbergen, 2008; Nicholls, 2009; Arruda and Hotamisligil, 2015). Under these conditions, dysfunction is associated with the mitochondrial permeability transition, a loss of inner mitochondrial membrane impermeability promoted by Ca^2+^ overload, oxidative imbalance and protein missfolding (Figueira et al., 2013; Biasutto et al., 2016; Vercesi et al., 2018).

We have recently found that mitochondria isolated from animals maintained on a chronic calorically‐restricted diet present increased Ca^2+^ accumulation capacity and resistance against mitochondrial permeability transition (Amigo et al., 2017; Menezes‐Filho et al., 2017), whereas acute fasting promotes increased susceptibility to permeability transition and reduced Ca^2+^ accumulation capacity (Menezes‐Filho et al., 2019). This further supports the notion that dietary interventions may not only affect physiological Ca^2+^ handling but also modulate damaging effects of supra‐physiological Ca^2+^ accumulation. Interestingly, caloric restriction and nutrient starvation also modulate mitochondrial morphology, stimulating mitochondrial fusion (Rambold et al., 2011; Khraiwesh et al., 2014; Cerqueira et al., 2016), while nutrient overload is often associated with mitochondrial fission (Molina et al., 2009; Liesa and Shirihai, 2013; Alsabeeh et al., 2018).

Altogether, a wealth of evidence supports links between nutritional status, mitochondrial morphology and dynamics, Ca^2+^ signaling and bioenergetic efficiency. However, a specific and central point that has not been studied is if changes in mitochondrial morphology directly promote changes in mitochondrial Ca^2+^ uptake that impact on intracellular Ca^2+^ signaling. Here we show that modifying mitochondrial morphology alters Ca^2+^ uptake rates and capacity, with larger mitochondria exhibiting faster and larger Ca^2+^ uptake. We also found that mitochondrial morphology regulates the cellular calcium‐replenishing process of store‐operated calcium entry (SOCE). Our results uncover a novel role for mitochondrial morphology in the control of mitochondrial Ca^2+^ uptake, which impact on cellular Ca^2+^ homeostasis.

## Methods

### Cell cultures

C2C12 cells (passages 6‐20) were cultured in high glucose DMEM with 10% FBS, 1 mM pyruvate, 100 units/mL penicillin and 1000 µg/mL streptomycin, trypsinized every 2‐3 days and typically split 1:7 (Wikstrom et al., 2012).

### Modulation of mitochondrial morphology

Mitochondrial fusion was promoted by infection with an adenovirus containing a dominant negative form of DRP1 (DRP1 DN, from Welgen, Inc.) at a multiplicity of infection level of 200. Mitochondrial fission was promoted by MFN2 silencing using an adenovirus from Welgen at a multiplicity of infection of 20. Cells were treated when split, and the media was changed after 24 hours. Experiments were conducted 72 hours after, when infection rates were maximized (Forni et al., 2016). Cell viability was assessed by trypan blue exclusion, and > 95% for all experiments conducted.

### Analysis of mitochondrial morphology

Cells were plated on glass‐bottomed plates 72 hours prior to imaging and loaded with Mitotracker Green (200 nM) for 30 min followed by wash‐out just prior to the experiment. A Zeiss LSM 880 confocal microscope in Airyscan mode was used for super‐resolution imaging, with a 488 nm Argon laser and Zeiss 63×/1.4NA oil immersion objective. At least 10 images per experimental condition were collected. Cells were individualized as areas of interest using Image J, and automated mitochondrial circularity and aspect ratios (the proportional relationship between width and height) were measured. Data are presented as symbols that represent average circularity and aspect ratio per cell. Mitochondrial fission is expected to increase circularity and decrease the aspect ratio. Representative images shown were adjusted in brightness and contrast for better visualization.

### Digitonin‐permeabilized cells

Cells were trypsinized, washed and suspended in 200 μL of 140 mM NaCl, 3 mM KCl, 400 μM KH_2_PO_4_, 20 mM Hepes, 5 mM NaHPO_4_, 5 mM glucose and 1 mM MgCl_2_, pH 7 (NaOH). They were counted and kept in this media at room temperature for at most 90 min while experiments were conducted, and ressuspended in permeabilized cell media (see below) just prior to each trace. Ideal cell quantities and digitonin titers were determined by following plasma membrane permeabilization using safranin (Kowaltowski et al., 2002) and were found to be 2.5 ⋅ 10^6^/mL cells in the presence of 0.0025% digitonin (added just prior to the trace). Under these conditions, cells were safranin‐permeant (indicating plasma membrane permeabilization) and maintained inner membrane potentials for 40 min (indicating mitochondrial membrane integrity).

### Mitochondrial Ca^2+^ uptake

Extramitochondrial Ca^2+^ uptake was followed in digitonin‐permeabilized cells using the florescent calcium probe Ca^2+^ Green (100 nM, Bambrick et al., 2006; Amigo et al., 2017) in media containing 125 mM KCl, 2 mM KH_2_PO_4_, 10 mM Hepes, 1 mM MgCl_2_, 5 mM succinate, 5 mM malate and 5 mM glutamate, pH 7 (KOH). Fluorescence was measured with constant stirring, in a cuvette fluorimeter at 506 nm excitation and 532 nm emission. Where indicated in the figures, successive additions of 50 μM CaCl_2_ were made, until Ca^2+^ accumulation capacity was exhausted, as indicated by lack of further uptake and/or release. A calibration curve was constructed using known CaCl_2_ concentrations, and maximal calcium uptake capacity and initial uptake rates were calculated for each trace. Where indicated, 1 μM cyclosporin A (CsA), a mitochondrial permeability transition inhibitor, or 1 μM ruthenium red (RR), an MCU inhibitor, were present. In Fig. 3, cells were treated either with 10 µM M1 1 or equivalent volumes of DMSO (control group) in the culture media 16 hours prior to membrane permeabilization and Ca^2+^ uptake measurements.

### Mitochondrial membrane potentials (ΔΨ)

Inner mitochondrial membrane potentials were determined in permeabilized cells by following the fluorescence of 5 μM safranin O at 485 nm excitation and 586 nm emission. For ideal digitonin permeabilization determination, the same media as Ca^2+^ uptake measurements was used, and the digitonin concentration that promoted permeabilization (seen as a low, stable, fluorescence) in less than 5 min, and sustained for at least 40 min was used. For ΔΨ quantification, media devoid of K^+^ was prepared (all K^+^ salts were substituted for Na^+^, except KCl, which was substituted by 250 mM sucrose). Fluorescence was related to known ΔΨs by adding 50 nM valinomycin and known KCl concentrations, allowing for ΔΨ to be calculated using the Nernst equation. A calibration curve was constructed relating ΔΨ and fluorescence, and extrapolated for basal fluorescence measured in the absence of added KCl (Kowaltowski et al., 2002).

### Ca^2+^ uptake in permeabilized cells with clamped ΔΨ

Cells were incubated in the same media used for ΔΨ quantification, with 50 nM valinomycin plus 0.41 mM KCl, sufficient to reach the same ΔΨ as found in MFN2 KD cells. Ca^2+^ uptake was measured as described above. Controls were conducted without the addition of KCl, in the same media. Samples were paired for preparations from the same flask/passage.

### Cellular Ca^2+^ imaging

Three days prior to the experiment, cells were plated in Greiner 627871 4‐compartment glass bottom cell culture dishes and mitochondrial morphology was modulated as described above. Fura2‐AM loading and imaging protocols were adapted from Arruda et al. (2017), with modifications. On the experimental day, the culture media was removed and cells were washed twice with experimental buffer containing 10 mM HEPES, 150 mM NaCl, 4 mM KCl, 1 mM MgCl_2_ and 10 mM D‐glucose (pH 7.4) and then incubated with 10 µM Fura2‐AM, 0.1% pluronic acid and 2 mM CaCl_2_ in that same media for 40 minutes at 37°C and 5% CO_2_ in order to load the probe. After incubation, media was removed and cells were washed twice in experimental buffer. Cells were then incubated in 1 mL experimental buffer and placed inside the chamber of a Leica DMi‐8 microscope equipped with a Fura2 filter (Leica Microsystems) and a 40× objective. Fura2 cytosolic Ca^2+^ imaging was conducted by alternatively illuminating the cells with wavelengths of 340 and 387 nm. Images were acquired every 5 s. Cytosolic Ca^2+^ levels are shown in the figures as the 340/387 ratios for the Fura2 channels, which are proportional to Ca^2+^ concentrations. Intensities were calculated using FIJI Image J extension by converting the videos obtained in each channel to grayscale and then plotting the mean gray values over time for each cell. Thirty seconds after the beginning of the experiment, 2 µM thapsigargin was added to the experimental media in order to promote ER Ca^2+^ release. At 630 s, 2 mM CaCl_2_ was added in order to trigger store‐operated Ca^2+^ re‐entry. Experiments were repeated on three separate days, with 7‐10 cells analyzed per day under each experimental condition. Cytosolic Ca^2+^ (340/387 rates) increase rates after the addition of 2 mM CaCl_2_ (Fig 2C) were measured by adjusting a linear fit using Origin 8 Pro software.

### ER stress marker evaluation by real‐time PCR

On the third day after infection, total RNA from the cells was extracted and purified using Trizol reagent (Invitrogen Life Technologies) and quality‐checked using 260/230nm and 260/280nm scores in a NanoDrop spectrophotometer. Equivalent contents of RNA were reverse‐transcribed using a High‐Capacity cDNA Reverse Transcription Kit (Thermo Fisher Scientific). Synthesized cDNA was stored at ‐20°C prior to the real‐time PCR assay. Data shown here are expressed as the ratio of the target gene to the HPRT reference/housekeeping gene, which was validated through deletion experiments (Nicot et al., 2005). cDNA amplification was performed using Platinum® SYBR® Green qPCR SuperMix UDG (Invitrogen Life Technologies) and evaluated by real‐time PCR using Rotor Gene 3000 apparatus (Corbett Research). qRT‐PCR primer sequences were obtained from the Harvard database PrimerBank (Wang et al., 2012), as listed in Supplementary Methods, and had their efficiency (90% minimum) and concentration standardized. Fold changes were calculated by the 2−ΔΔCT method.

### Statistical analysis

Comparisons were made using Graphpad Prism software, with t‐tests for simple comparisons between two groups and ANOVA for multiple comparisons.

## Results

To study the effects of mitochondrial morphology modulation on Ca^2+^ uptake by this organelle we used C2C12 myoblast cells, which have a functional and dynamic mitochondrial network (Wikstrom et al., 2012; Sin et al., 2016), are amenable to automated mitochondrial morphological quantification (Valente et al., 2017), and display robust intracellular Ca^2+^ signaling (Gutierrez‐Martin et al., 2005). Figure 1 (leftmost panels) shows typical images of stained mitochondria in these cells, such as those used for automated morphological analysis. We found that DRP1 DN cells, in which the fission machinery is inhibited, presented mitochondria with lower circularity (Fig. 1A) and higher aspect ratios (the proportional relationship between width and height, Fig. 1B) when compared to control cells. This indicates that DRP1 DN cells, as expected, had longer and less circular mitochondria as a consequence of the inhibition of fission. Conversely, MFN2 KD cells, in which fusion was partially impaired (mRNA levels were decreased on average by 40%, results not shown), presented enhanced mitochondrial fragmentation, as indicated by higher circularity (Fig. 1A) and lower aspect ratios (Fig. 1B). Notably, the changes in morphology were expected for physiological conditions (Liesa and Shirihai, 2013). Neither genetic interference affected the area occupied by mitochondria (Fig. 1C), indicating no large‐scale disruption of mitochondrial content occurred. Furthermore, cell viability was above 95% under all conditions, showing that we were able to successfully modulate mitochondrial morphology without compromising cell integrity.

**Figure 1:**
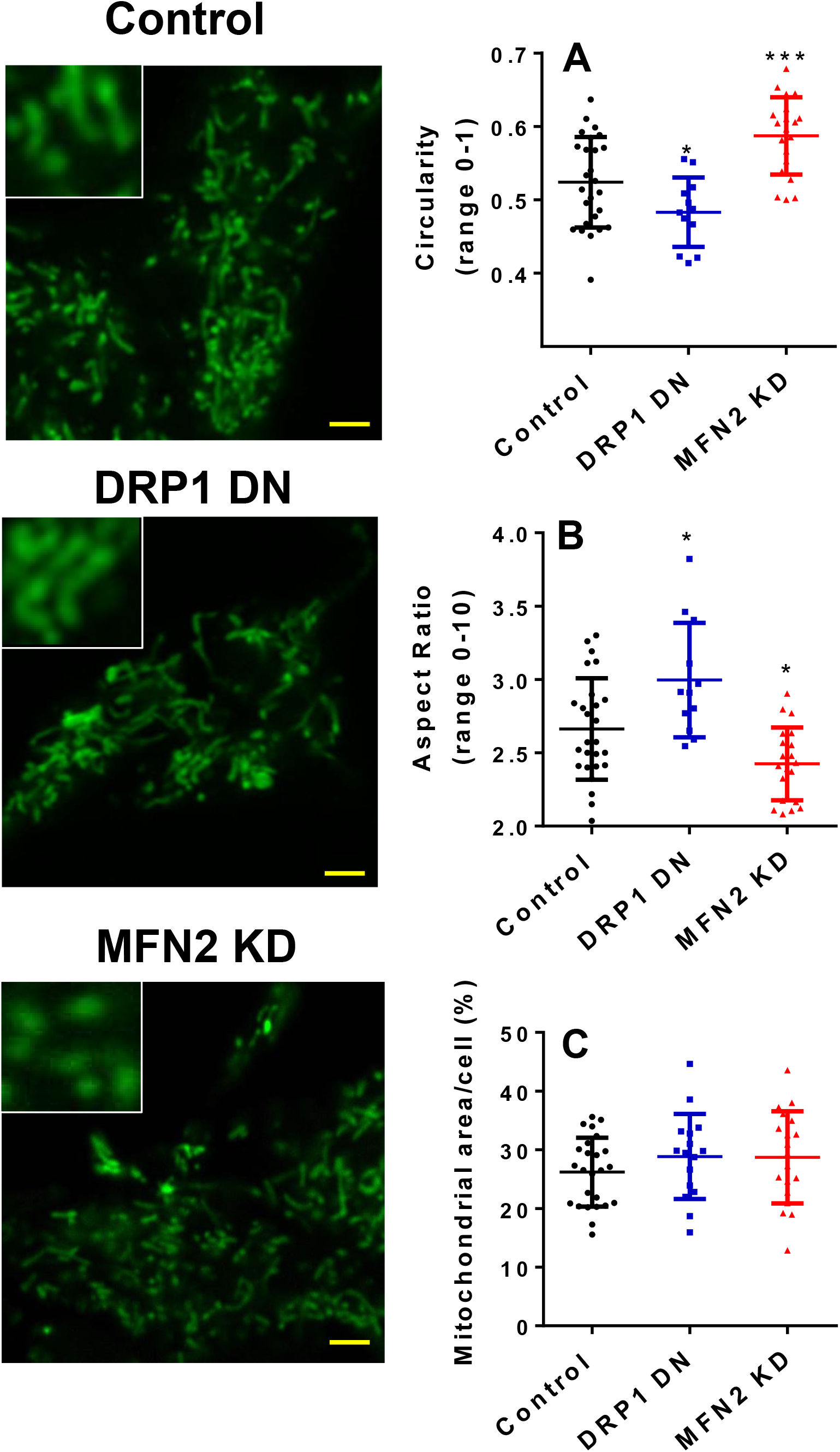
Mitochondrial morphology is modulated by DRP1 and MFN2. C2C12 cell mitochondria were marked with MitoTracker Green and imaged in intact cells as described in the Methods section (Left Panels, scale bar = 10 μm). The inserts show higher magnifications of selected areas. Mitochondrial circularity (Panel A), aspect ratios (Panel B) and cross‐sectional area relative to the cellular area (Panel C) were quantified from these images. N > 10 cells per condition, *p < 0.05, ***p < 0.001 relative to control cells.

Next, we determined if the changes in mitochondrial morphology modified Ca^2+^ uptake by these organelles using digitonin‐permeabilized cell suspensions. Plasma membrane permeabilization under these conditions has been shown to keep mitochondrial and cell architecture, as well as the cytoskeleton, intact (Fiskum et al., 1980), while largely diluting cytosolic components, allowing for a direct assessment of organellar Ca^2+^ dynamics. Calcium levels were followed in permeabilized cells using the membrane‐impermeable probe Calcium Green, which fluoresces in the presence of Ca^2+^ in the experimental buffer, but does not enter membrane‐bound organelles. Fig. 2A shows a scheme of the experimental setup. Adding successive Ca^2+^ boluses (where indicated by the arrows) to a suspension of permeabilized cells results in a rapid increase in Ca^2+^ levels, followed by gradual decrease in Ca^2+^ detection due to the uptake of the ion by membrane‐bound organelles. Uptake of the ion continues after each bolus addition until the maximal capacity is reached. Beyond this point, addition of Ca^2+^ to the extramitochondrial media is not followed by its uptake by these organelles. Eventually, calcium overload lead to spontaneous Ca^2+^ release.

**Figure 2:**
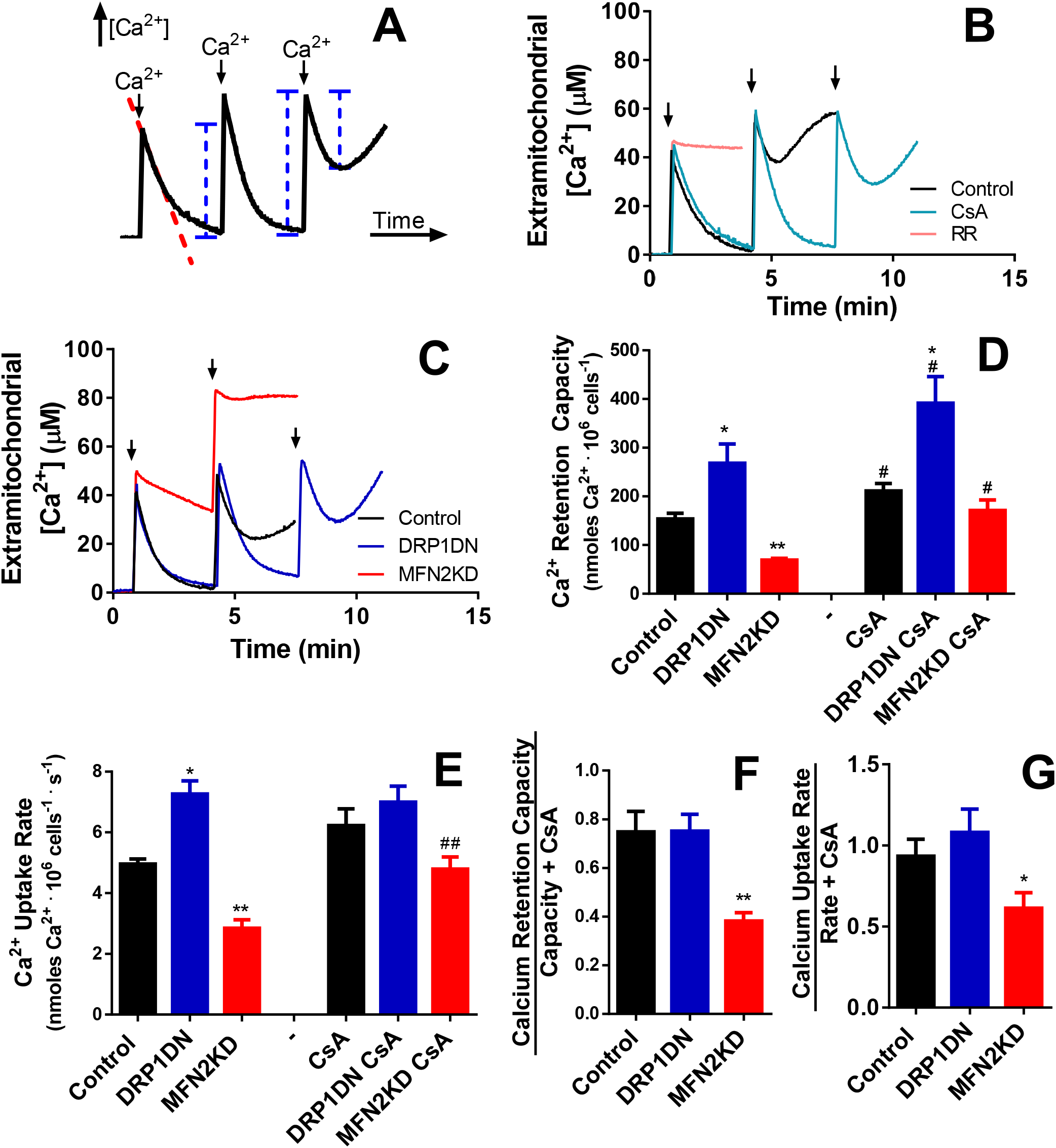
Ca^2+^ uptake is modulated by mitochondrial morphology. Ca^2+^ uptake was measured in permeabilized cells as described in the Methods section, in control (black lines/columns), DRP1 DN (Panels B‐D; dark blue lines/columns) and MFN2 KD (Panels B‐D, red lines/columns) cells, as indicated. Panel A indicates the measurement technique used: Ca^2+^ uptake by mitochondria is followed by measuring extramitochondrial [Ca^2+^] concentrations over time, after the addition of successive boluses of the ion to the experimental media. The dotted red line indicates data used to measure Ca^2+^ uptake rates. Blue lines represent Ca^2+^ quantities that, when added, reflect maximal retention capacity. In Panel B, only control cells were used, and, where indicated, 1 μM CsA (light blue line) or 1 μM RR (pink line) were present. Arrows indicate bolus 50 μM Ca^2+^ additions. Panel C shows typical traces in control (black), DRP1 DN (blue) and MFN2 KD (red) cell types. Arrows indicate bolus 50 μM Ca^2+^ additions. Panel D quantifies Ca^2+^ retention capacity in traces such as those shown in Panel C, while Panel E quantifies Ca^2+^ uptake rates over the first 20 sec after the first Ca^2+^ addition in traces such as those in Panel C. Basal calcium retention capacity relative to retention in the presence of CsA (Panel F) and basal uptake rates relative to rates in the presence of CsA (Panel G) were plotted. N ≥ 5 independent repetitions, *p < 0.05, **p < 0.01 relative to non‐infected cells under the same conditions. ^#^p < 0.05, ^##^p < 0.01, relative to the same cells in the absence of CsA.

Ca^2+^ removal from the media is due to mitochondrial activity, as uptake by this organelle occurs with higher capacity than Ca^2+^ uptake by the ER. Indeed, Ca^2+^ uptake in control cells (Fig. 2B, black line) was completely prevented in the presence of the mitochondrial calcium uniporter (MCU) inhibitor ruthenium red (RR, Fig. 2B, pink line). Inhibition of the mitochondrial permeability transition with cyclosporin A (CsA, Fig. 2B, light blue line) resulted in a large increase in Ca^2+^ uptake capacity, further confirming that mitochondrial Ca^2+^ uptake was measured.

Two characteristics of mitochondrial Ca^2+^ uptake can be calculated from traces generated by this assay: Calcium retention capacity, the maximal amount of Ca^2+^ taken up by mitochondria (see blue lines in Fig. 2A) before spontaneous release, and calcium uptake rates (see the dotted red line in Fig. 2A), the speed in which the ion is removed from the media after the first addition. We found that DPR DN cells, in which mitochondria become elongated, have higher calcium retention capacity (blue traces in Fig. 2C, and blue columns in Fig. 2D) and higher calcium uptake rates (Fig. 2E) when compared to control cells (black traces and columns, Figs 2C‐E). On the other hand, MFN2 KD cells, with fragmented mitochondria, present lower calcium uptake capacity (Fig. 2C, red trace and Fig. 2D, red columns) and lower Ca^2+^ uptake rates (Fig. 2E, red columns). Overall, this shows that increased mitochondrial fusion enhances Ca^2+^ uptake speed and capacity, while mitochondrial fission decreases Ca^2+^ uptake. On the other hand, the affinity for Ca^2+^ in the cells with different mitochondrial morphologies was equal, as evaluated by residual concentrations of the ion left in the media after uptake stabilized.

The presence of cyclosporin A (CsA, Fig. 2D) enhanced calcium retention capacity in all cell types, although uptake with cyclosporin A was still larger in DRP1 DN mitochondria and lower in MFN2 KD. Interestingly, CsA had a particularly pronounced effect on MFN2 KD mitochondrial Ca^2+^ uptake. We calculated the proportional effect of the mitochondrial permeability transition on Ca^2+^ uptake capacity and rates, by comparing these values in the absence and presence of CsA. Our results indicate that proportional susceptibility to permeability transition is not changed in DRP1 DN cells, but this process is facilitated by mitochondrial fission promoted by MFN2 knockdown: both mitochondrial Ca^2+^ uptake capacity (Fig. 2F) and rates (Fig. 2G) are proportionally lower in MFN2 KD cells when normalized to these measurements in the presence of CsA. Thus, MFN2 KD‐promoted fragmentation increases mitochondrial permeability transition activity. However, despite the effects of permeability transition in MFN2 KD cells, the absolute differences in Ca^2+^ uptake capacity and rates promoted by changes in mitochondrial morphology persisted even in the presence of CsA (Fig. 2D and E). This shows that the permeability transition is not solely responsible for the changes in Ca^2+^ uptake observed with modified mitochondrial morphology, but rather an additional factor governing Ca^2+^ homeostasis in this organelle.

Changes in MCU expression have been described in cells derived from MFN2 knockout animals, but do not occur in acute MFN2 knockdown cells such as our model (Filardi et al., 2015). In our experiments, changes in MCU expression could explain altered uptake rates, but could not account for the changes in calcium retention capacity seen. Regulation of the MCU also does not seem to be involved in the modulation of Ca^2+^ uptake by mitochondrial morphology, since spermine, an MCU activator, did not enhance mitochondrial uptake in control or MFN2 KD cells (not shown). In order to verify if an independent and more acute morphology‐modulating intervention could reproduce the effects seen by genetically manipulating mitochondria, we tested the effects of fusion‐promoting compounds. In preliminary experiments, mdivi‐1, identified as a mitochondrial division inhibitor in a chemical screen (Cassidy‐Stone et al., 2008), promoted extensive cell death within 16 hours, possibly because of its effect as a Complex I inhibitor (Bordt et al., 2017). A second small molecule, mitochondrial fusion promoter M1, did not affect cell viability when tested, and was used to induce acute fusion in the cells (Fig. 3). M1 treatment robustly increased both calcium retention capacity (Fig. 3A) and uptake rates (Fig. 3B) in control cells, confirming through a pharmacological approach that mitochondrial fusion promotes an increase in Ca^2+^ uptake by this organelle.

**Figure 3:**
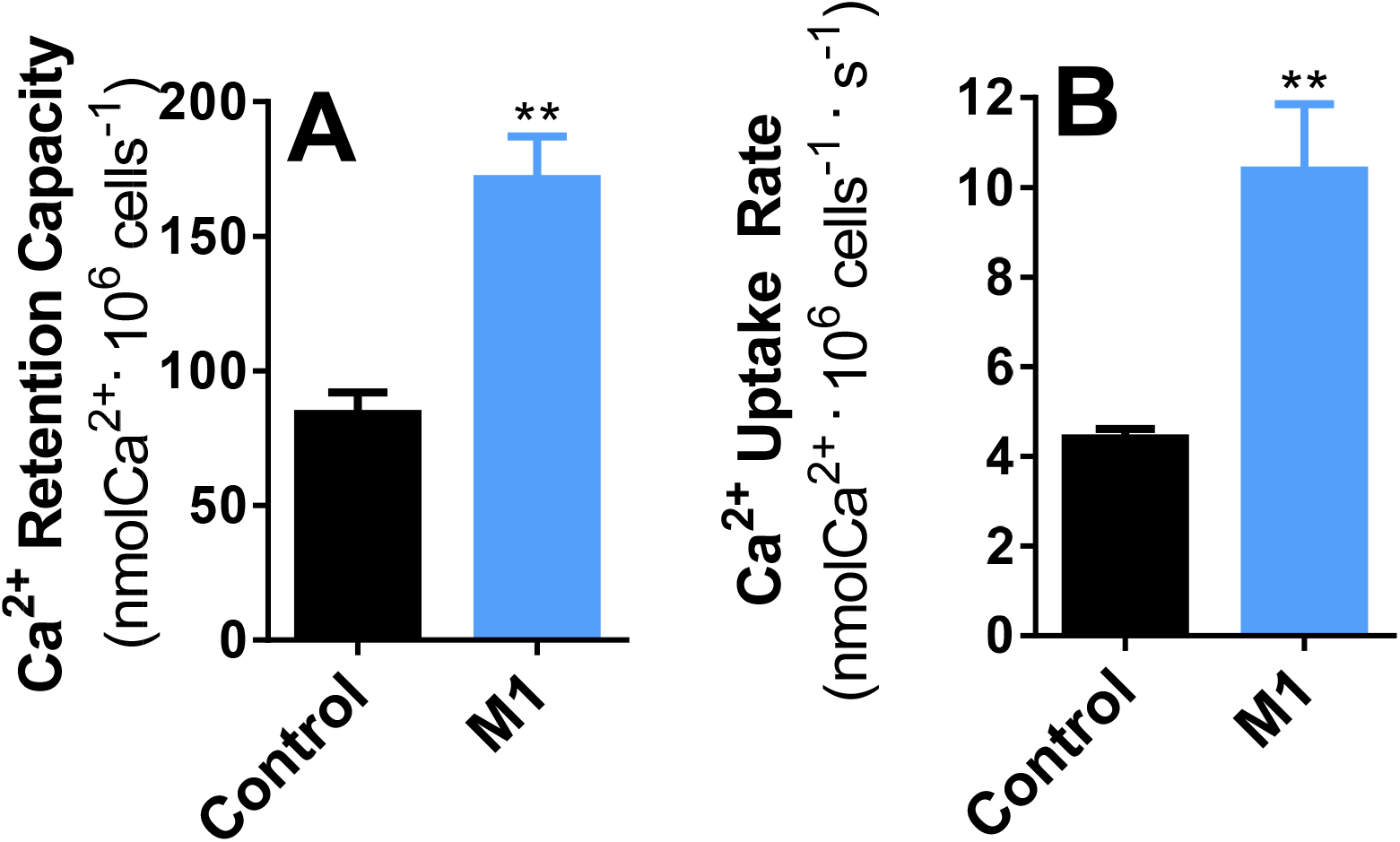
Mitochondrial fusion promoted by M1 increases Ca^2+^ uptake. Control cells were incubated in the presence or absence of 10 µM M1 as described in the Methods session, and Ca^2+^ retention capacity (Panel A) and uptake rates (Panel B) were measured as described for Fig. 2. N = 4 independent repetitions, **p < 0.01 relative to DMSO‐treated (control) cells.

The fact that mitochondrial Ca^2+^ homeostasis could be modulated by two different methods to change mitochondrial morphology (genetically manipulating protein levels and use of a fusion‐inducing small molecule) led us to seek a mechanistic explanation for the changes in Ca^2+^ uptake. A possible reason for the differences in Ca^2+^ uptake observed could be changes in mitochondrial inner membrane potentials (ΔΨ), the driving force for Ca^2+^ uptake. We thus measured mitochondrial ΔΨ using plasma membrane‐impermeable safranin O fluorescence, which allows for quantitative ΔΨ determinations in isolated mitochondria or permeabilized cells (Figure 4; Akerman and Wikström, 1976; Kowaltowski et al. 2002). Fig. 4A shows typical safranin fluorescence traces over time after cell permeabilization with digitonin. The downward deflection of the curve over time is a result of ΔΨ‐driven accumulation of the cationic probe in the mitochondrial matrix, which decreases its fluorescence (Akerman and Wikström, 1976; Kowaltowski et al., 2002). Safranin accumulation in DRP1 DN cells (blue trace) was consistently higher than in control cells (black trace), while MFN2 KD cells (red trace) accumulated less safranin. This result could be indicative of changes in ΔΨ promoted by manipulating mitochondrial morphology. However, accumulation and fluorescence of ΔΨ‐sensitive probes is also modified by mitochondrial size and shape (Kowaltowski et al., 2002; Kowaltowski, 2019). To overcome this possible artifact and quantify changes in ΔΨ, we calibrated safranin fluorescence curves in a K^+^‐free media in the presence of the K^+^ ionophore valinomycin, by adding known quantities of extramitochondrial K^+^ into the media and following fluorescence changes (Fig. 4B; Akerman and Wikström, 1976). Under these conditions, ΔΨ can be calculated for each added K^+^ concentration using the Nernst equation, and basal ΔΨ can be extrapolated from the calibration curves, in millivolts (Akerman and Wikström, 1976; Kowaltowski et al. 2002). FCCP, a proton ionophore, was added at the end of each trace to complete the dissipation of ΔΨ. By using K^+^ calibrations, we found that, despite differences in safranin accumulation and fluorescence, control and DRP1 DN mitochondria displayed the same ΔΨ (in the range of 180 mV, Fig. 4C), while MFN2 KD mitochondria had significantly lower ΔΨ (in the range of 155 mV, Fig. 4C). Despite this lower ΔΨ, MFN2 KD mitochondria were still capable of promoting oxidative phosphorylation, as demonstrated by measuring a decrease in ΔΨ upon the addition of 2 mM ADP (result not shown). These results highlight a highly important and often overlooked caveat of studies using mitochondrially‐accumulated probes under conditions in which morphology is modified: In the absence of calibration for each individual condition, changes in fluorescence may be misinterpreted as changes in the parameter the probe is measuring (Kowaltowski et al., 2002; Kowaltowski, 2019), when they may only reflect variations in calibration.

**Figure 4:**
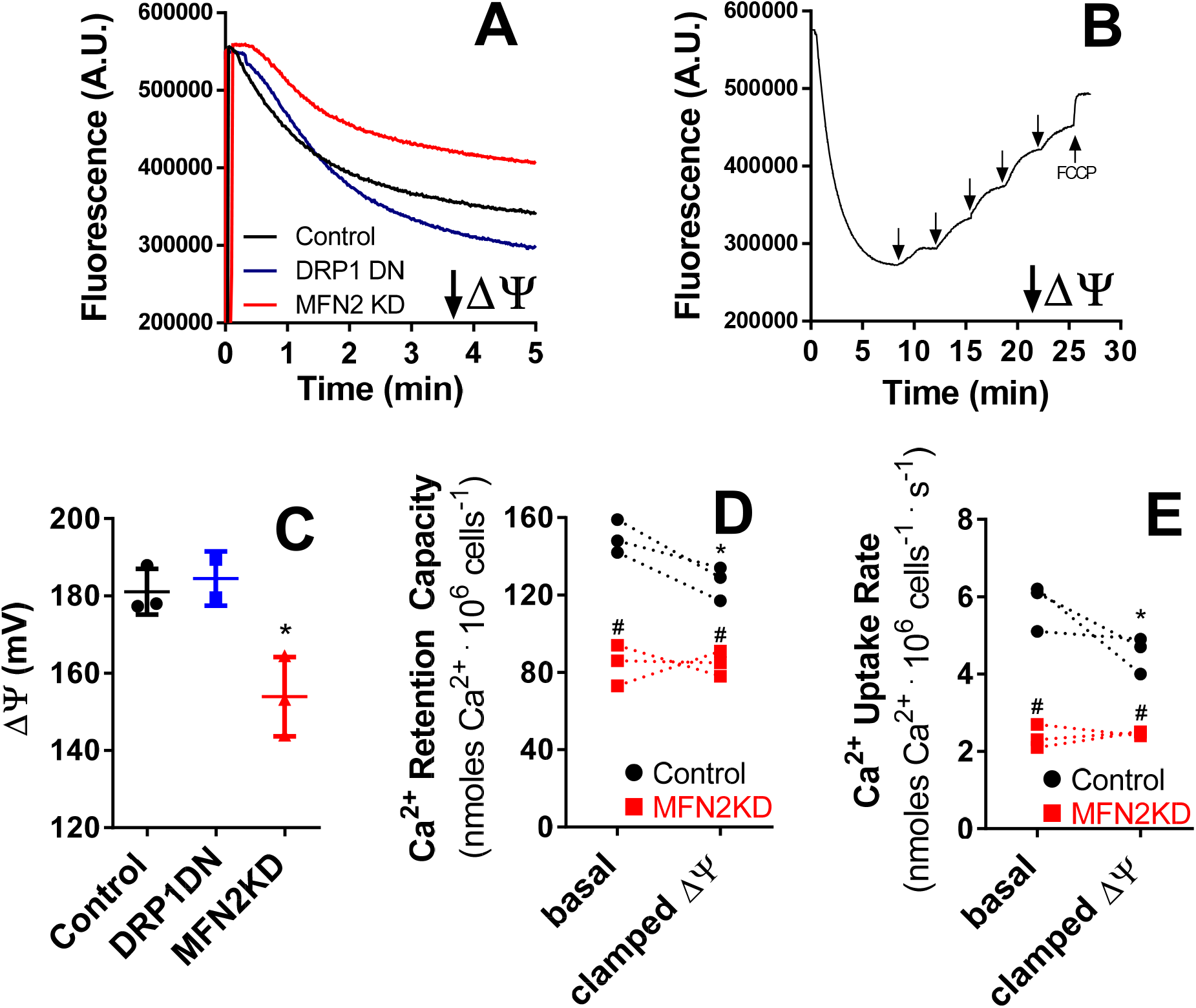
Mitochondrial morphology affects inner mitochondrial membrane potential (ΔΨ) measurements. Mitochondrial inner membrane potentials (ΔΨ) were measured using safranin O fluorescence as described in the Methods section. Panel A shows typical safranin fluorescence traces in control (black), DRP1 DN (blue) and MFN2 KD (red) permeabilized cells. Panel B shows a typical ΔΨ calibration procedure in control cells, conducted as described in the Methods section. Arrows indicate where K^+^ and the uncoupler FCCP (1 μM) were added. Panel C presents calibrated ΔΨ determinations in control (black symbols), DRP1 DN (blue symbols) and MFN2 KD (red symbols) permeabilized cells. n = 5, *p < 0.05 relative to control cells. In Panel D, Ca^2+^ uptake capacity was evaluated in control (black circles) and MFN2 KD (red squares) permeabilized cells under basal conditions or when ΔΨ was clamped to the average measured MFN2 KD ΔΨ (see Methods). In Panel E, Ca^2+^ uptake rates were evaluated in control (black circles) and MFN2 KD (red squares) permeabilized cells under basal conditions or when ΔΨ was clamped to the average measured MFN2 KD ΔΨ. N = 3 independent repetitions, *p < 0.05 relative to basal ΔΨ. ^#^p < 0.05 relative to control cells under the same ΔΨ condition; traces connect paired experiments.

Overall, our ΔΨ quantifications do not explain the increase in Ca^2+^ uptake capacity and rates in DRP1 DN mitochondria, since ΔΨ was unchanged relative to control cells. However, the decrease in ΔΨ observed in MFN2 KD mitochondria may be the cause for lower Ca^2+^ uptake capacity and rates as well as higher permeability transition susceptibility observed in Fig. 2. To assess this possibility, we designed an experiment in which the ΔΨ of control cell mitochondria was forcibly clamped at the same level as the ΔΨ of MFN2 KD mitochondria using added extramitochondrial K^+^ and valinomycin (see Methods). Calcium uptake was then quantified under these conditions, comparing the effects of clamping ΔΨ on Ca^2+^ retention capacity (Fig. 4D) and uptake rates (Fig. 4E). MFN2 KD cells presented equal calcium retention and uptake rates under basal and clamped ΔΨ conditions (red squares; connected traces show individual replicates under basal conditions paired with the same sample with clamped ΔΨ). This result was expected, since ΔΨ was clamped at the same level as basal ΔΨ in MFN2 KD cells. On the other hand, control cells (black circles) presented a decrease in retention capacity and uptake rates when the ΔΨ was clamped. However, mitochondrial Ca^2+^ uptake capacity and rates were still significantly higher in control cells relative to MFN2 KD cells under clamped ΔΨ conditions, indicating that ΔΨ is not the only factor decreasing Ca^2+^ uptake in cells with impaired mitochondrial fusion. This result is in line with the finding that DRP1 DN mitochondria do not present changes in ΔΨ, but take up more Ca^2+^, at faster rates. Furthermore, changes in ΔΨ would be expected to impact upon uptake rates, but not necessarily retention capacity. Overall, our results conclusively demonstrate that changes in mitochondrial morphology and dynamics are sufficient to change Ca^2+^ homeostasis in this organelle.

Our next question was if these modifications in mitochondrial Ca^2+^ homeostasis have an impact on cellular calcium handling. To investigate this, we evaluated cytosolic Ca^2+^ levels in intact cells in which mitochondrial morphology was modulated (Figure 5). Fig. 5A shows typical Fura 2 fluorescence ratio traces, which are directly proportional to intracellular calcium concentrations. We found that MFN2 KD cells had consistently lower basal calcium levels (typical traces in Fig. 5A are quantified in Fig. 5B), while levels in control and DRP1 DN cells were equal. Upon promoting calcium release from the ER with thapsigargin (Thapsi, as indicated by the arrow in Fig. 5A), MFN2 KD cells showed lower cytosolic Ca^2+^ increments (Fig. 5B) relative to control and DRP1 DN cells, indicative of lower ER Ca^2+^ stores. After ER calcium release, basal cytosolic Ca^2+^ levels were re‐quantified, and once again found to be lower in MFN2 KD cells (Basal 2, Fig. 5B). Overall, these results show that MFN2 KD cells had lower cytosolic and ER Ca^2+^ concentrations.

**Figure 5:**
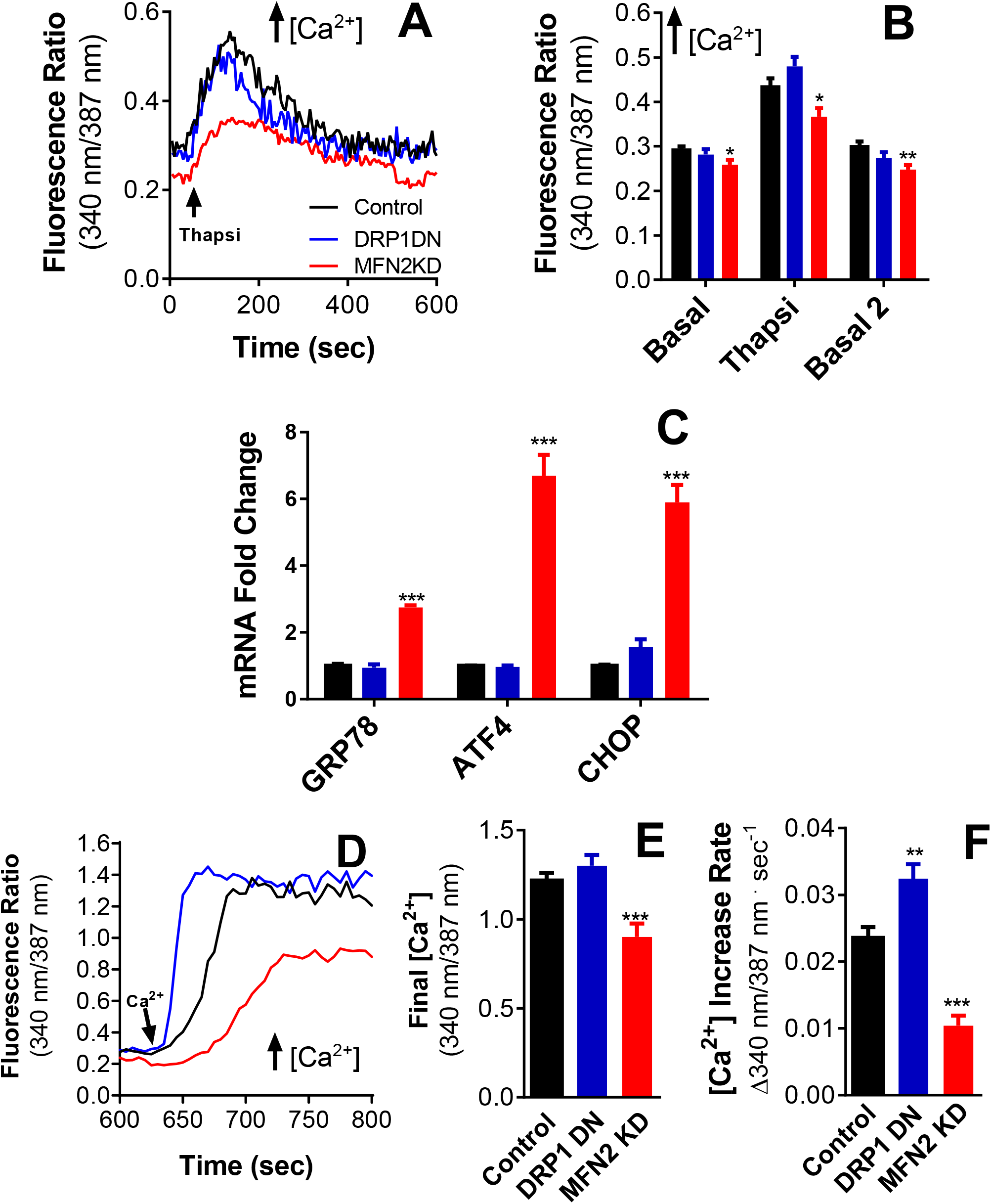
Mitochondrial morphology affects cellular Ca^2+^ homeostasis and ER stress responses. Panel A shows representative traces of Fura2 ratios corresponding to cytosolic Ca^2+^ concentrations in intact control (black), DRP1 DN (blue) and MFN2 KD (red) plated cells. Where indicated, 2 µM thapsigargin was added. Panel B shows quantifications of these traces, including basal initial ratios, peak thapsigargin ratios and “basal 2” ratios, after thapsigargin. In Panel C, mRNA levels of ER stress markers GRP78, ATF4 and CHOP are expressed as the fold change relative to the housekeeping gene HPRT. Panel D shows representative traces of Fura2 ratios after the addition of 2 mM extracellular Ca^2+^, to activate SOCE. Final SOCE fluorescence ratios are quantified in Panel E. Panel F quantifies SOCE fluorescence increase rates. N = 23‐27 cells per condition, *p < 0.05, **p < 0.01, ***p < 0.001 compared to control cells.

Disruption of ER Ca^2+^ homeostasis is linked to ER stress responses (Krebs et al., 2015), so we evaluated the expression levels of ER stress markers GRP78 (an ER chaperone also known as BiP; Lee, 2005), ATF4 and CHOP (transcription factors that induce protein synthesis in the unfolded protein response; Han et al., 2013) in our cells (Fig. 5C) as a second indication of a disruption in ER Ca^2+^ homeostasis. We found that the mRNA expression of all three proteins was strongly upregulated in MFN2 KD cells compared to control or DRP1 DN cells, indicating that the low basal and ER Ca^2+^ levels in MFN2 KD cells were accompanied by ER stress.

Since ER calcium stores were depleted in MFN2 KD cells, we additionally evaluated store‐operated calcium entry (SOCE), or the capacity to activate Ca^2+^ influx into the cell from the extracellular environment as a mechanism to compensate the emptying of intracellular Ca^2+^ stores (Parekh and Putney, 2005). SOCE is known to be regulated by mitochondria, through mechanisms that involve active mitochondrial Ca^2+^ uptake and release (Malli and Graier, 2017; Spät and Szanda, 2017; Ben‐Kasus et al., 2017). We measured SOCE in our cells by re‐adding extracellular Ca^2+^ to cells preincubated in Ca^2+^‐ free media and in which ER stores had been previously depleted by thapsigargin. This promoted a rapid re‐entry of the ion into the cytosol, increasing Fura 2 fluorescence ratios (Fig. 5D). We found that SOCE into MFN2 KD cells was impaired, reaching lower maximal calcium when compared to control cells (Fig. 5E) and occurring at slower rates (Fig. 5F). This result is compatible with the depletion of ER Ca^2+^ stores we found previously. DRP1 DN cells, on the other hand, displayed maximal calcium replenishment similar to control cells (Fig. 5E), but at significantly faster rates (~30% faster, Fig. 5F). Taken together, these experiments demonstrate that regulating mitochondrial morphology has an expressive impact on different aspects of cellular physiological Ca^2+^ handling.

## Discussion

We demonstrate here that inducing moderate changes in the morphology of the mitochondrial network, promoting either fission or fusion (Fig. 1), alters mitochondrial Ca^2+^ uptake and retention properties as well as cellular Ca^2+^ homeostasis and ER stress. Specifically, promoting mitochondrial fission through MFN2 KD enhances mitochondrial permeability transition (Fig. 2), possibly due to its effects of decreasing ΔΨ (Fig. 4), a known inducer of this process (see Vercesi et al., 2018 for a review). MFN2 KD also significantly changes mitochondrial Ca^2+^ homeostasis: it decreases both Ca^2+^ uptake rates and Ca^2+^ retention capacity in mitochondria (Fig. 2). Additionally, MFN2 KD has effects on cellular ion homeostasis, as it lowers basal Ca^2+^ levels and ER Ca^2+^ stores, activates ER stress and hampers store‐operated Ca^2+^ entry (SOCE) after intracellular Ca^2+^ store depletion (Fig. 5). While these last effects seen in intact cells may be related to MFN2’s properties in mediating the interaction between mitochondria and the ER (Pernas and Scorrano, 2016), it is important to note that mitochondrial Ca^2+^ uptake assays conducted in permeabilized cells are independent of these interactions, since Ca^2+^ is added directly to the extramitochondrial microenvironment, and uptake by the ER is not quantitatively relevant (Fig 2B).

Experiments promoting mitochondrial fusion suggest that the Ca^2+^ uptake effects seen in MFN2 KD cells are related to mitochondrial morphology itself. Mitochondrial fusion in DRP1 DN cells (Fig. 2) or M1‐treated (fission‐inhibited) cells (Fig. 3) significantly enhances both mitochondrial Ca^2+^ uptake capacity and rates, in a manner independent of changes in ΔΨ (Fig. 4). Although mitochondrial permeability transition may be responsible for the increments in uptake rates in DRP1 DN cells relative to control cells (uptake rates are equal in these cells in the presence of CsA, Fig. 2E), it does not account for enhanced Ca^2+^ uptake capacity, which remains higher in more fused mitochondria even in the presence of a permeability transition inhibitor (Fig. 2D). Indeed, enhanced Ca^2+^ uptake is probably attributable to mitochondrial morphological changes themselves, since these change organellar matrix capacity, where the ions accumulate. Demonstrating that mitochondrial fusion also impacts on cellular Ca^2+^ physiology, DRP1 DN cells had similar basal Ca^2+^ levels and ER Ca^2+^ stores, but displayed enhanced SOCE rates relative to control cells (Fig. 5D,F).

This finding complements prior work showing that mitochondrial function, and specifically Ca^2+^ uptake and release, is determinant in SOCE‐mediated cellular Ca^2+^ store replenishing (Malli and Graier, 2017; Spät and Szanda, 2017). Importantly, it suggests that mitochondrial morphology may be an important physiological regulator of SOCE, which was to date been related to mitochondrial function through the use of non‐physiological stimuli such as uncouplers or direct inhibition of Ca^2+^ uptake or release pathways in these organelles (Malli and Graier, 2017; Spät and Szanda, 2017; Ben‐Kasus et al., 2017). Interestingly, there is also evidence that the location of mitochondria near plasma membrane Ca^2+^ entry sites can be important in SOCE (Fonteriz et al., 2016), a result compatible with our results, since the regulation of mitochondrial morphology also impacts on mitochondrial transport and positioning in the cell (Anesti and Scorrano, 2006).

In a more global sense, our results demonstrate a tight link between mitochondrial morphology, cellular and mitochondrial Ca^2+^ homeostasis. These findings are supported by prior experiments that indirectly suggest an association between mitochondrial morphology and Ca^2+^ uptake into this organelle. For example, Szabadkai et al. (2004) showed that DRP1‐mediated mitochondrial fission disrupted mitochondrial networks and impacted on intraorganellar calcium wave propagation, indicating a role for mitochondrial plasticity in cell‐wide mitochondrial Ca^2+^ dissemination. Additionally, Lewis and co‐workers (2018) recently found that MFF‐mediated mitochondrial fission changed axonal Ca^2+^ homeostasis and impacted on synaptic function due to changes in mitochondrial Ca^2+^ uptake, connecting this effect to mitochondrial mass in the axon. Both results can also be explained by a decrease in mitochondrial Ca^2+^ uptake rates and capacity secondary to mitochondrial fission, as seen in our present study. Our results are also in line with the work of Sebastián et al. (2012), who found that MFN2 knockout mice presented elevated levels of ER stress markers, similar to our findings in MFN2 KD cells, although the authors did not connect these findings to Ca^2+^ changes.

Many central important biological events involve simultaneous changes in mitochondrial morphology and in Ca^2+^ homeostasis, including immune activation (Vig and Kinet, 2009; Baixauli et al., 2011), differentiation (Tonelli et al., 2012; Forni et al., 2016), insulin secretion (Flatt et al., 1980; Stiles and Shirihai, 2012) and fatty acid metabolism (Otto and Ontko, 1978; Rambold et al., 2015; Benador et al., 2019), among others. It is tempting to speculate that at least part of the regulatory mechanisms in these processes involve changes in mitochondrial and cellular Ca^2+^ homeostasis promoted by the modulation of mitochondrial morphology we describe here.

## Acknowledgements

This work was funded by *Fundação de Amparo à Pesquisa do Estado de São Paulo* (FAPESP) CEPID grant number 2013/07937‐8, CAPES (Coordenação de Aperfeiçoamento de Pessoal de Nível Superior Finance Code 001), *Conselho Nacional de Pesquisa e Desenvolvimento* (CNPq), NIH‐NIDDK 5‐R01DK099618‐02 and RO1DK56690, UCLA Department of Medicine Chair commitment and UCSD/UCLA Diabetes Research Center pilot grant, NIH P30 DK063491. The funders had no role in study design, data collection and analysis, decision to publish, or preparation of the manuscript. Edson Alves Gomes and Camille Caldeira da Silva are acknowledged for outstanding technical support. P.N. is supported by FAPESP fellowship number 2014/24511‐7. S.L.M.F. was a CAPES PhD fellowship recipient.

## Abbreviations

CsA: cyclosporin A
DMEM: Dulbecco’s modified Eagle medium
DRP1: dynaminrelated protein 1
ER: endoplasmic reticulum
FBS: fetal bovine serum
MCU: mitochondrial calcium uniporter
MFN2: mitofusin 2
RR: ruthenium red
SOCE: storeoperated calcium entry
ΔΨ: mitochondrial inner membrane potential

