## Supplementary methods for "Mitochondrial Morphology Regulates Organellar Ca^2+^ Uptake and Changes Cellular Ca^2+^ Homeostasis"

Primers used from Thermo Fisher

| **Protein** | **Foward sequence** | **Reverse sequence** |
|---|---|---|
| ***Atf4*** | GAGCTTCCTGAACAGCGAAGTG | TGGCCACCTCCAGATAGTCATC |
| **BiP/GRP78** | TTCAGCCAATTATCAGCAAACTCT | TTTTCTGATGTATCCTCTTCACCAGT |
| **CHOP** | CCACCACACCTGAAAGCAGAA | AGGTGAAAGGCAGGGACTCA |

Primers used from Exxtend (obtained using the Harvard database PrimerBank)

| **Protein** | **Foward sequence** | **Reverse sequence** |
|---|---|---|
| **Mfn2** | AGAACTGGACCCGGTTACCA | CACTTCGCTGATACCCCTGA |
| **HPRT** | TCAGTCAACGGGGGACATAAA | GGGGCTGTACTGCTTAACCAG |
